# Toward a dynamic threshold for quality-score distortion in reference-based alignment

**DOI:** 10.1101/754614

**Authors:** Ana A. Hernandez-Lopez, Claudio Alberti, Marco Mattavelli

## Abstract

The intrinsic high entropy sequence metadata, known as quality scores, are largely the cause of the substantial size of sequence data files. Yet, there is no consensus on a viable reduction of the resolution of the quality score scale, arguably because of collateral side effects. In this paper we leverage on the penalty functions of HISAT2 aligner to rebin the quality score scale in such a way as to avoid any impact on sequence alignment, identifying alongside a distortion threshold for “safe” quality score representation. We tested our findings on whole-genome and RNA-seq data, and contrasted the results with three methods for lossy compression of the quality scores.

## Introduction

High-throughput sequencing technologies epitomize the challenges of data-intensive science, the current paradigm for scientific exploration (*1*). Massive datasets, diverse in variety, exhaustive in scope, and fine-grained in resolution are representative descriptors of sequence data, the big data (*2*) in the biomedical sciences. The drop in sequencing costs (*3*) has democratized the production of sequence data and propelled its growth. Sequence data has historically doubled every seven months ^a^, a figure that is meticulously tracked and updated by the GenBank, the genetic sequence database of the National Institute of Health (NIH), every two months. The current storage requirements for sequence data are, conservatively, on par with the storage estimates for any of the other major Big Data producers: Astronomy (1 EB/year), Twitter (0.001-0.017 EB/year) and YouTube (1-2 EB/year) (*4*). It has been projected an annual storage need between 2-40 EB per year for sequence data. However, we are at the onset of the sequence data flood. The promise of personalized medicine to revolutionize the diagnosis and treatment of diseases has triggered projects to sequence an important proportion of the human population. An attest to this is the 100 000 Genomes Project, a study launched in the UK that sequenced one hundred thousand human genomes from patients with rare diseases and their families, and patients with cancer (*5*). As per estimates of The Global Alliance for Genomics and Health, more than 60 million patients will have their genome sequenced by 2025 (*6*), a projection that will be facilitated by the competition of private companies to offer genome sequencing services at a population scale with milestones to reduce sequencing costs, currently pushing the cost of 100 dollars per sequenced genome. The ubiquitous integration of personal genomic information into aspects of everyday life is around the corner, as we step into era of the “social genome” (*7*). The capacity to generate massive datasets of sequence data greatly outpaces our ability to analyze them, the notorious bottleneck in omic analyses. As per projections to the year 2025, over 75% of the cost and complexity in omic workflows will be taken over by data analysis and storage (*8*). Data is accumulating very rapidly, and the immediate, pressing challenge in high-throughput sequencing is to reduce the size of sequence data files, the FASTQ files, for storage. Many solutions in the realm of data compression have emerged to reduce the size of such files, and much work has been devoted to explore methods to compress sequence data, and its associated metadata, without loss of information (*9, 10*).

Substantial storage size is dedicated to sequence metadata in lossless compressed files compared to the storage devoted to sequence data. Conservatively speaking, this figure is over 50% in lossless compressed files. There has been early evidence in the scientific literature to the size occupied by sequence metadata (*11, 12*), and more recently such proportions have been brought back to attention (*13, 14*). Sequence metadata, also referred to as quality scores, is the bottleneck in the compression of FASTQ files.

The study of lossy quality score representation was initiated as a remedy to the large storage footprints of FASTQ files. The investigation of techniques for lossy compression of quality scores, along with quantification of their impact on the calling of genetic variants has been well studied. It has been shown and confirmed at length that lossy approaches for quality score representation provide significant storage saving with negligible impact on variant calling (*11, 13, 15–18*), regardless the idiosyncrasies of the lossy compression approach. The effect of lossy compression of quality scores has also been explored in differential gene expression with similar conclusions on the negligible effect of applying lossy representations (*19*). Furthermore, recent advances in sequencing technologies are leading the production of longer genomic sequences with better accuracy and drastically reduced resolution for the quality scores (*20*), supporting the claim that coarser representations are in principle suitable for omic analyses.

While the ultimate interest lies in assessing the impact of lossy quality score representation in a full bioinformatic pipeline, this approach may be ineffective for the purpose of understanding the role of lossy quality scores in the analysis. This is because sequence data is transformed continually as it is shepherded throughout the pipeline, and errors and associated uncertainties of the computational methods that operate on it are combined along with the data, obscuring the precise effect of lossy quality scores in the analysis.

Moreover, despite several efforts to evidence marginal impact in the application of lossy quality score representation, no consensus exist on the limits of a “safe” representation for compressing them lossy. In this context, this work focuses on read alignment to explore the effect of lossy quality score representation. In particular, we show that it is possible to compute a threshold value for transparent quality score distortion for sequence alignment, allowing the identification of a safe representation for the quality score scale. A result that aligns with current trends in sequencing technologies pushing for coarser resolutions to reduce the storage footprint of sequence data.

### Read alignment

The challenge to represent lossy quality scores in the alignment of sequence reads lies in maintaining the reads original alignment location(s) with the new simplified representation. In quality-aware aligners, quality score values participate in the computation of suitable alignment locations for the reads. The way in which quality score values is factored in depends on the alignment technique, and their usage is not essential but clearly optional. Many aligners have been developed that do not rely on quality scores. This is readily noted in benchmark comparisons, which commonly include widely used aligners (*21–25*).

Using quality scores can improve alignment accuracy because the information they provide, the probability of error in the calling of each sequence base, can be incorporated to determine which positions in a read are more important to map (*21,26*). Quality scores can be used in very diverse ways among alignment tools, as the methods prioritize this metadata differently.

One of the most widely used reference-based aligners, BWA (*27, 28*), incorporates quality scores in a measure for the reliability of alignments. The aligner does this by defining a mapping quality score that represents the error probability of each read alignment. Quality scores are not used in BWA’s alignment algorithm but rather they are used to support alignment results. For example, alignments of high accuracy can be screen out using the mapping quality score. Moreover, this score is used by the aligner to estimate the insert size distribution in paired-end mapped reads.

In constrast, quality scores can be incorporated at the core of an aligners algorithm to guide the alignment decision. This is the case for Novoalign (*29*), another well-known reference-based aligner, which consistently ranks well in alignment accuracy. Novoalign uses quality score information in its penalization system to score candidate alignment locations for each input sequence read.

Our purpose is to investigate the contribution of quality scores to alignment in HISAT2, which uses quality score metadata for the computation of alignment scores. Built over Bowtie2 (*30*), HISAT2 is in fact the evolution of this very well-know and popular aligner. It has good adoption and performance (*23, 31*), and has stood the test of time. Moreover, it is open source, and it is still been maintained ^b^. In addition, it was designed to map both DNA and RNA-seq reads.

The role of quality scores in alignment is framed within HISATs scoring system, and understanding it will be the way through finding a simplified representation for the quality scores that circumvents undesirable effects on alignment. Concretely, the goal is to preserve alignment locations as if no modification to the values of quality scores was done before alignment, that is, we aim at varying the quality scores transparently.

We ask, under what circumstances quality score values are, or become, informative for determining the alignment location of a read? To address this question we look into how quality scores weigh in on HISAT2’s scoring system.

## Methods

Aligning sequences consists in lining up characters to reveal similarity. However, the aligner cannot always assign a read to its point of origin with high confidence, thus it makes an educated guess about its origin in the reference sequence.

HISAT2 quantifies how similar the sequence of a read is to the reference sequence it aligns to by computing an alignment score (AS) for the read, whose value is used by the aligner to classify reads as aligned or unaligned. Therefore, the AS can be seen as a proxy to measure the effect quality scores have on alignment.

The aligner starts with the assumption that no difference exists between the read sequence `r` and the segment of the reference sequence `R`, pointed to by the alignment location, it aligns to. If this condition is satisfied, the best possible alignment score is assigned to the read, which is zero. This the largest, non-negative value the alignment score can take. The concept of alignment score does not apply to unaligned reads, as such HISAT2 does not report a value, nor the AS metric for these reads. As dissimilarities are found between `r` and `R`, HISAT2 penalizes each discrepant sequence character. Penalty values are always negative and are added together to compute the total alignment score for the read `r`. For an alignment to be considered good enough, or valid, it must have an alignment score with a value no less than the minimum score threshold `t`. The threshold is configurable and a function of the read length x, and its default value is `t(x) = 0 − 0.2 × (x)`. Thus, valid alignments meet or exceed the minimum score theshold and are capped at zero. For example, aligned sequences of 100 base-pairs long will have valid alignment scores in the range `−20 ≤ AS ≤ 0`.

There are four types of penalizations, and each is scored differently:

- Ambiguous characters (N). The penalty is set in positions where the read, reference or both, contain an ambiguous character such as N. For each ambiguous character the penalization is 1
- Gaps. Affine gaps in the read or the reference are penalized for their occurrence (gap opening, `O`), and for each position they span (gap extension `E`). The sum of both values defines the penalization for the gap. The penalty for a read gap of length `n` is `O + n × E`, and its default is `5 + n × 3`. The same expression applies for a reference gap of length `n` Soft-clips (sc).
- Reads can be aligned in a way such that they are trimmed at one or both extremes, because some of the characters at their ends do not match the reference. Omitted characters are trimmed or soft-clipped from the read to produce a valid alignment. Each character that is soft-clipped receives a penalty value defined by the penalty function

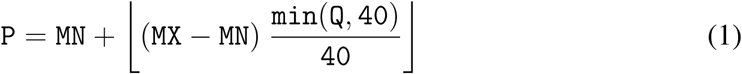

where `MX = 2`, and `MN = 1` are the default values, and `Q` is the quality score for the soft-clipped character
- Mismatches (mm). These are discrepant characters between the read and the reference. Each mismatch is penalized using the penalty function for soft-clips. However, the parameters values change for mismatches, and they default to `MX = 6`, and `MN = 2`

Lets note that quality scores participate only in the penalization for mismatches and soft-clips. By solving the penalty function above for the full quality score scale, and for both mismatches and soft-clips, we get Table 1.

**Table 1:** Penalty values for mismatches and soft-clips. Penalties are always negative values. P_mm_: Penalization for mismatches; P_sc_: Penalization for soft-clips.

| ASCII | Q | $f = \frac{\min(Q,40)}{40}$ | $P_{mm} = - ( 2 + \lfloor 4 \times f \rfloor )$ | $P_{sc} = - ( 1 + \lfloor 1 \times f \rfloor )$ |
| --- | --- | --- | --- | --- |
| 73 | 40 | 1 | -6 | -2 |
| 72 | 39 | 0.975 | -5 | -1 |
| 71 | 38 | 0.95 |  |  |
| 70 | 37 | 0.925 |  |  |
| 69 | 36 | 0.9 |  |  |
| 68 | 35 | 0.875 |  |  |
| 67 | 34 | 0.85 |  |  |
| 66 | 33 | 0.825 |  |  |
| 65 | 32 | 0.8 |  |  |
| 64 | 31 | 0.775 |  |  |
| 63 | 30 | 0.75 |  |  |
| 62 | 29 | 0.725 | -4 |  |
| 61 | 28 | 0.7 |  |  |
| 60 | 27 | 0.675 |  |  |
| 59 | 26 | 0.65 |  |  |
| 58 | 25 | 0.625 |  |  |
| 57 | 24 | 0.6 |  |  |
| 56 | 23 | 0.575 |  |  |
| 55 | 22 | 0.55 |  |  |
| 54 | 21 | 0.525 |  |  |
| 53 | 20 | 0.5 |  |  |
| 52 | 19 | 0.475 | -3 |  |
| 51 | 18 | 0.45 |  |  |
| 50 | 17 | 0.425 |  |  |
| 49 | 16 | 0.4 |  |  |
| 48 | 15 | 0.375 |  |  |
| 47 | 14 | 0.35 |  |  |
| 46 | 13 | 0.325 |  |  |
| 45 | 12 | 0.3 |  |  |
| 44 | 11 | 0.275 |  |  |
| 43 | 10 | 0.25 |  |  |
| 42 | 9 | 0.225 | -2 |  |
| 41 | 8 | 0.2 |  |  |
| 40 | 7 | 0.175 |  |  |
| 39 | 6 | 0.15 |  |  |
| 38 | 5 | 0.125 |  |  |
| 37 | 4 | 0.1 |  |  |
| 36 | 3 | 0.075 |  |  |
| 35 | 2 | 0.05 |  |  |
| 34 | 1 | 0.025 |  |  |
| 33 | 0 | 0 | 8 |  |

The hypothesis is that sequence alignment is preserved when quality score distortion and alignment score invariance occur simultaneously. To test this, we start by grouping the quality score scale shown in Table 1, according to the penalty values for mismatches and soft-clips. The result is the table shown in Figure 1.

**Figure 1:**
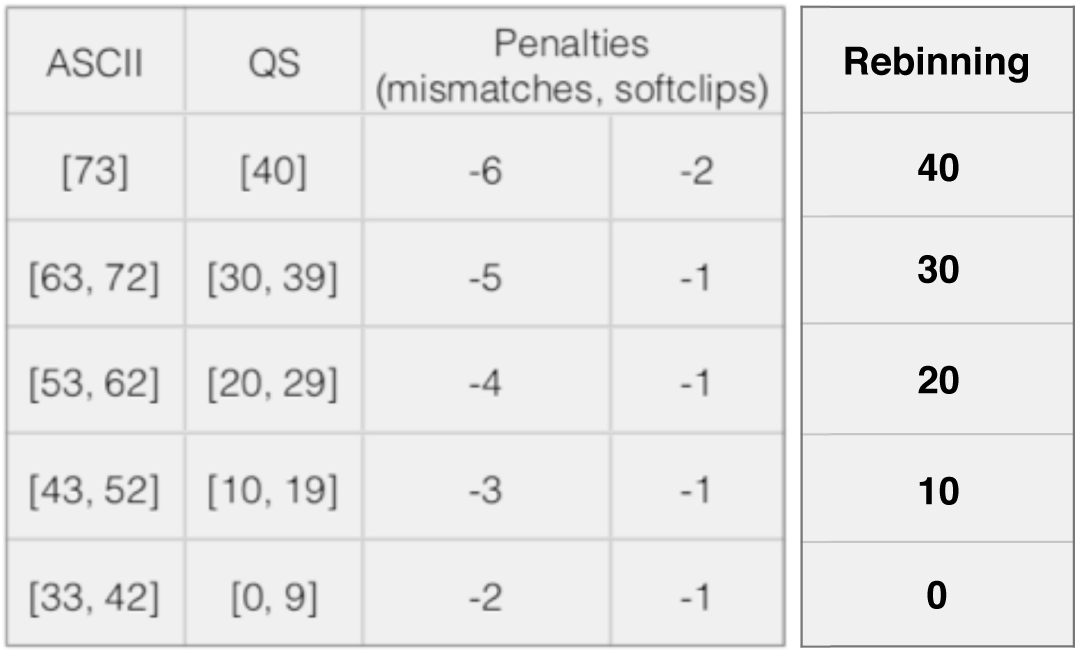
Rebinning of quality score scale.

With this rebinning we can compute distortion rate baselines that represent lossy compression rates that can “at least” be applied to the quality scores of raw sequence files (FASTQ files) without compromising alignment. These baselines can be thought of as distortion thresholds, which rely on sequence files. Figure 2 shows the setup of our experiments. An input file with undistorted quality scores (**D**) is rebinned to produce an output file with distortion rate **d**. Both undistorted and rebinned files are aligned, and produce identical alignment reports. The distortion threshold for file **D** is **d**.

**Figure 2:**
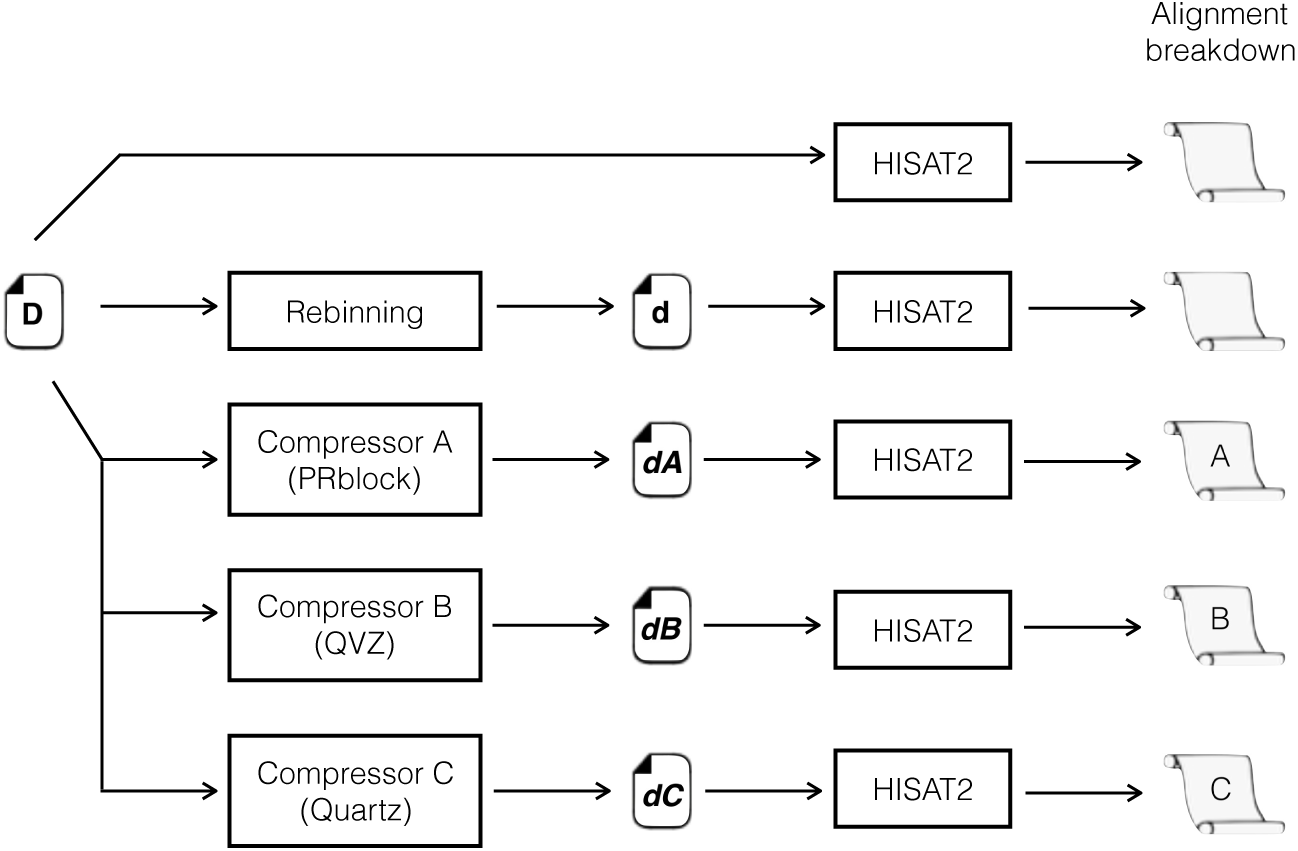
Experimentation setup.

To observe the effect that quality score distortion plays on alignment we ran three lossy compressors: PRblock (*32*), QVZ (*33*), and Quartz (*34*), and set their parameters such that the output files met as close as possible the value of the distortion threshold **d**. The approximate distortion rates for each compressor are ***dA, dB*** and ***dC*** (refer to Figure 2). The distorted files were then aligned with HISAT2 to quantify mapping results.

## Results

We experimented with synthetic and natural data and are reporting results for two natural data samples: T16M Metastatic liver tumor (whole-genome sequence data) (*35*), and Gene expression data in skin fibroblast cells (rna-seq data) (*36*). Results are reported in the tables in Figure 3. The alignment report is presented as the percentage of reads grouped in one of three possible sets: reads that aligned zero times (`Z`), reads that aligned exactly one time (`X`), and reads that aligned more that one time (`M`).

**Figure 3:**
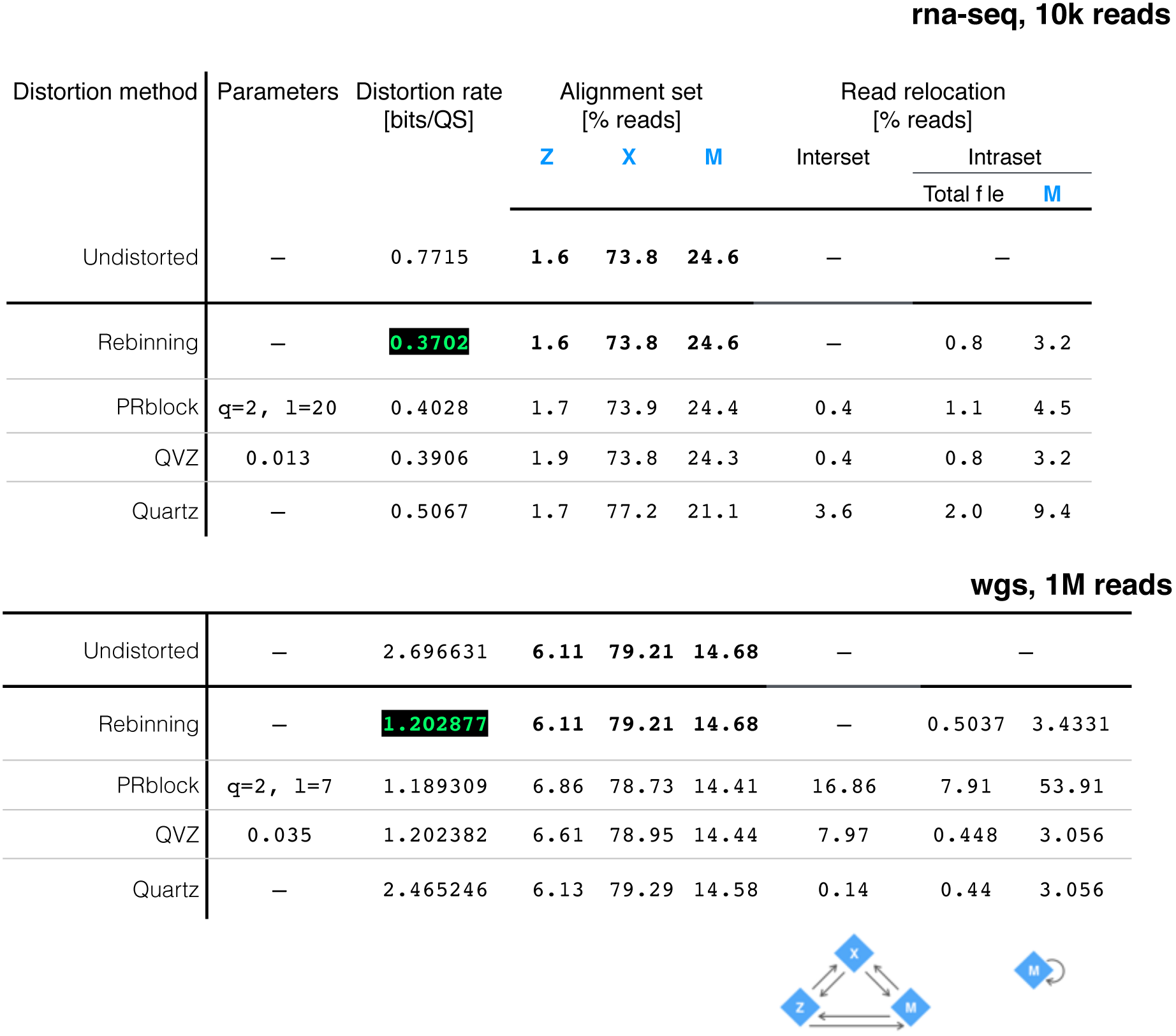
Distortion rate and alignment percentages for wgs and rna-seq samples.

The tables summarize alignment information as the percentage of reads whose alignment coordinate changed as a consequence of quality score distortion. We call this read relocation, and can happen between alignment sets or within alignment set `M` (see Figure 4).

**Figure 4:**
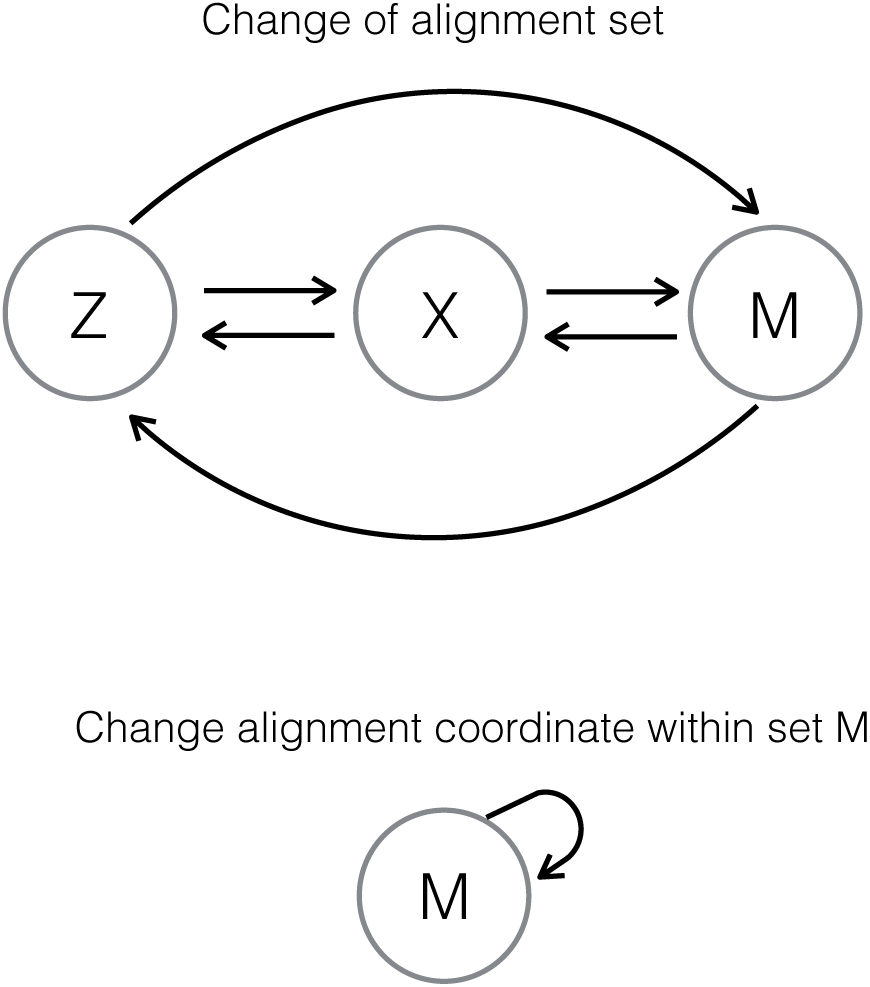
Read relocation between sets (top), and within set `M` (bottom).

For example, a read aligned before quality score distortion may be grouped in set `Z` but if that same reads is aligned after quality score distortion it may be grouped in set `X`. This type of read relocation is between sets, or interset, and the percentage of reads relocated in this fashion is shown under Interset read relocation in Figure 3.

The second form of read relocation can occur within set `M`, when the quality scores of a read with multiple alignment locations are modified in a way such that the new alignment coordinate belongs to the set of its multiple candidate locations. The percentage of reads relocated within set `M` is shown under Intraset read relocation in Figure 3. The percentages shown are relative to the total file and to the set of multireads (M).

Note that this type of read relocation occurs even in the rebinned file. This happens when the set `M` contains reads whose set of alignment coordinates have the same alignment score. HISAT2 will select one of the candidate coordinates for each read by computing a pseudo-random number generated from the read name, the sequence string, the quality score string and an optional seed value. Thus, modifying the quality scores will trigger HISAT2 intrinsic response toward multireads with equally likely alignment coordinates.

The graphs in Figure 5 report the effect of rebinning. In both graphs, the points to the far right show the lossless compression rate for each file. In this case, no changes to the quality scores are made, and therefore, all alignment coordinates are preserved.

**Figure 5:**
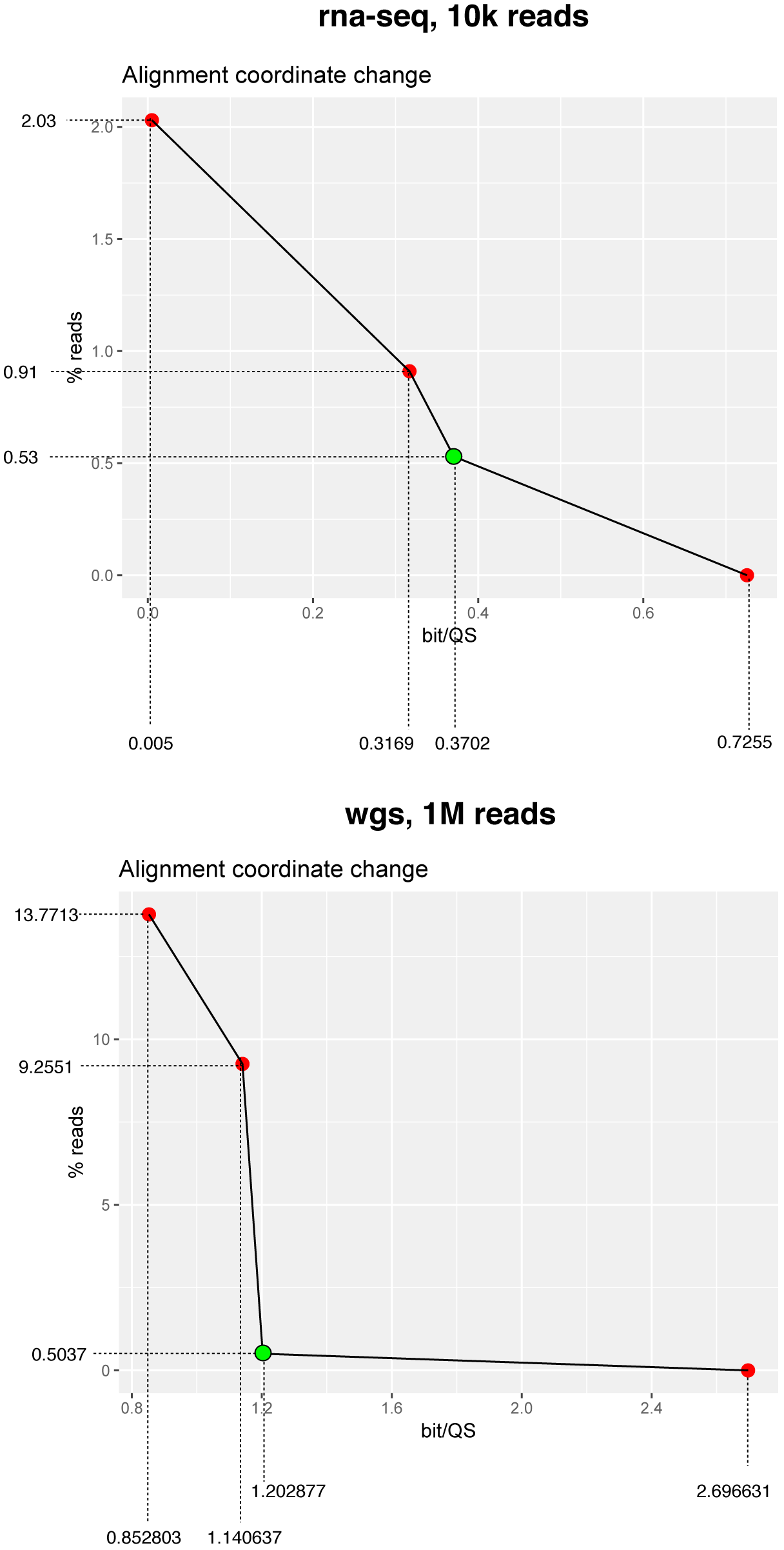
Alignment coordinate changes for rna-seq and wgs samples

If we then rebin the quality score scale by changing their values according to Figure 1, in five bins, the file can be compressed at a rate indicated by the green dots in both graphs in Figure 5. The green dots identify distortion thresholds for AS invariance, and every rebinned file has a specific threshold value. Notice there is a percenage of alignment coordinate changes, which result as a consequence of intraset read relocation of multireads. In Figure 5, the red points to the left of the green dots show the rebin of the quality score scale but this time using three and two bins, instead of the standard five bins shown in Figure 1. Notice the abrupt raise in the percentage of affected reads as we move toward the left of the graph, a consequence of pushing for a coarser representation for the quality score scale.

Changes to the quality scores of read sequences will inevitably lead to changes in alignment coordinates, therefore impacting alignment. Assessing the significance of this impact will depend on the recipient application following sequence alignment. However, the impact of lossy quality scores on alignment can be eliminated by keeping the alignment scores invariant. Although this is in principle true, we discovered that some idiosyncratic design decisions in the aligner weigh in unexpectedly, and collaterally impact alignment locations; this is beyond our control.

Sequence reads will fall in one of thee sets after alignment, and an alignment location(s) will be reported afterward for each read. As seen at the top of Figure 6, each read `U` will be assigned an alignment score `AS` by the aligner, and this value will determine whether the read receives an alignment position or not. Aligned reads are grouped by the number of locations the aligner found for them; if that number is one, they group in set `X`, if more than one location are found for a read, they group in set `M`. An unaligned read has no alignment location and belong to set `Z`.

**Figure 6:**
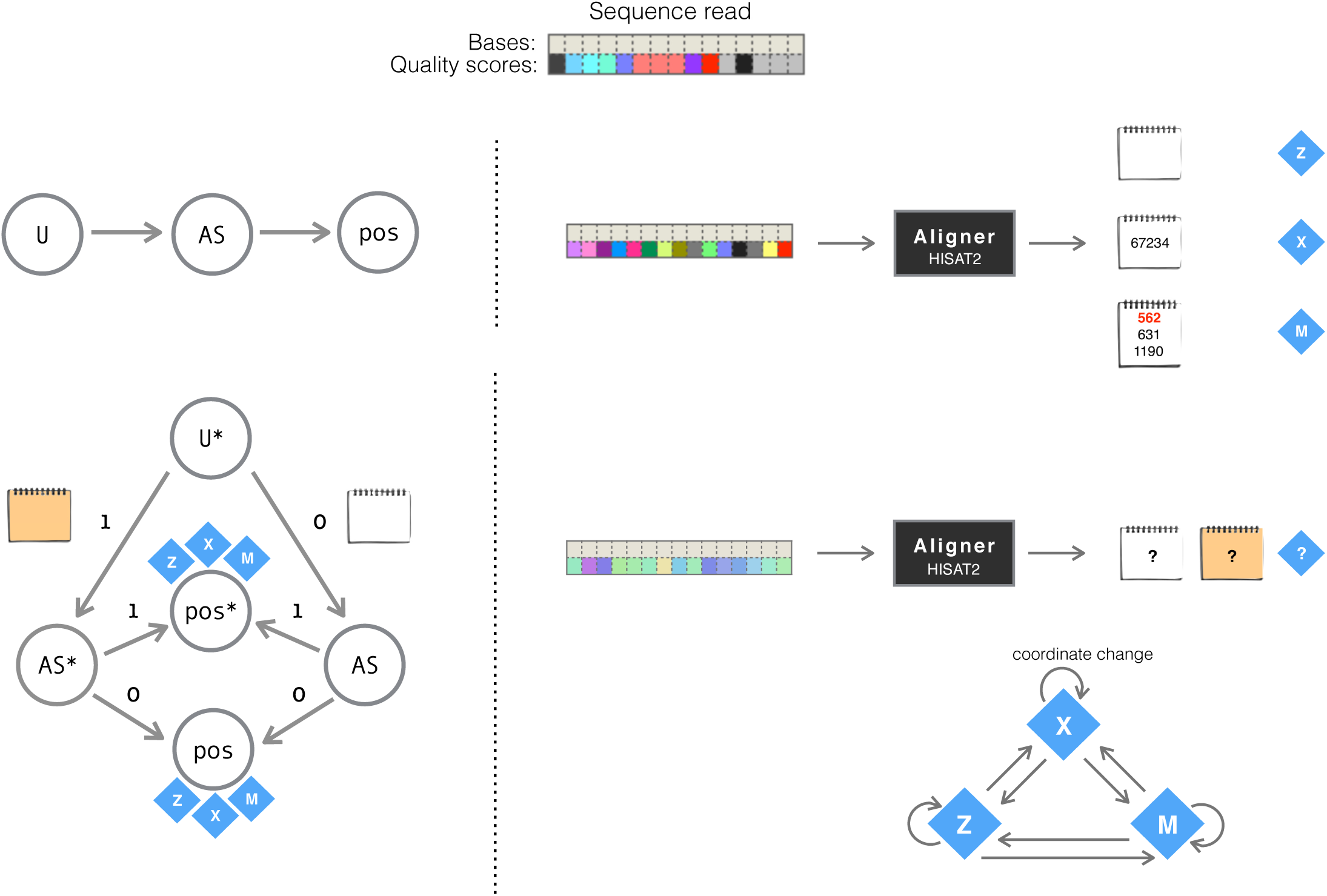
Abstraction of the assignment of alignment location(s) for each input read. In the figure, alignment coordinates are depicted by the notepads. A read aligned as a multiread is reported with list of possible alignment locations, and one of them is randomly selected as the primary alignment (shown in red).

When changes are made to their quality scores of a read `U`*, and it is then aligned, the report of its alignment score may change (`AS`*), or keep the same value (`AS`). Refer to the bottom of Figure 6. A change in the value of the AS does not immediately yield a change in the alignment position for that read. However, it could be the case that it does (`pos`*), and thus the alignment coordinate is tracked to record its displacement. Regardless of the outcome, the read will group in either `Z`, `X` or `M` set.

The effect of rebinning reads, in accordance to Figure 1, and aligning them afterward is shown in Figure 7. Invariance of alignment scores is achieved for every input read `U`*, and the only reads that could potentially be affected by this new representation to their quality scores are the multireads. Therefore, changes to alignment coordinates can happen only within set `M`.

**Figure 7:**
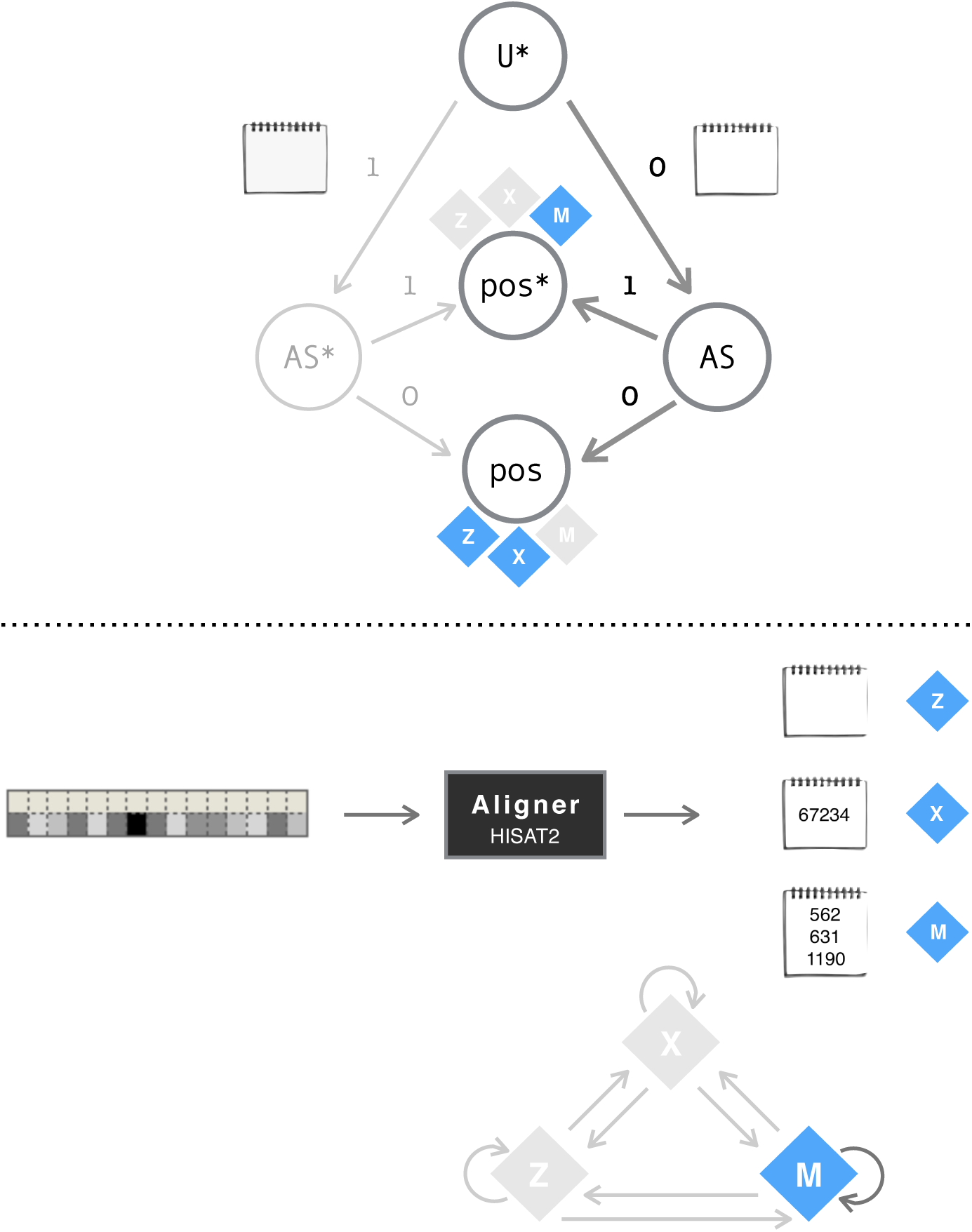
Abstraction of the assignment of alignment location(s) for rebinned reads. For multi reads, the report of the primary alignment is randomly selected by the aligner. No guarantees can therefore be given with respect to their values.

A graphical summary of the process is presented in Figure 8. Input reads without changes, and with changes (rebinning), to their quality scores are aligned and compared side to side. The content of the three alignment sets (`Z`, `X` and `M`) are preserved. As for the alignment coordinates, they are kept unchanged with no guarantees for those in the mutiread set.

**Figure 8:**
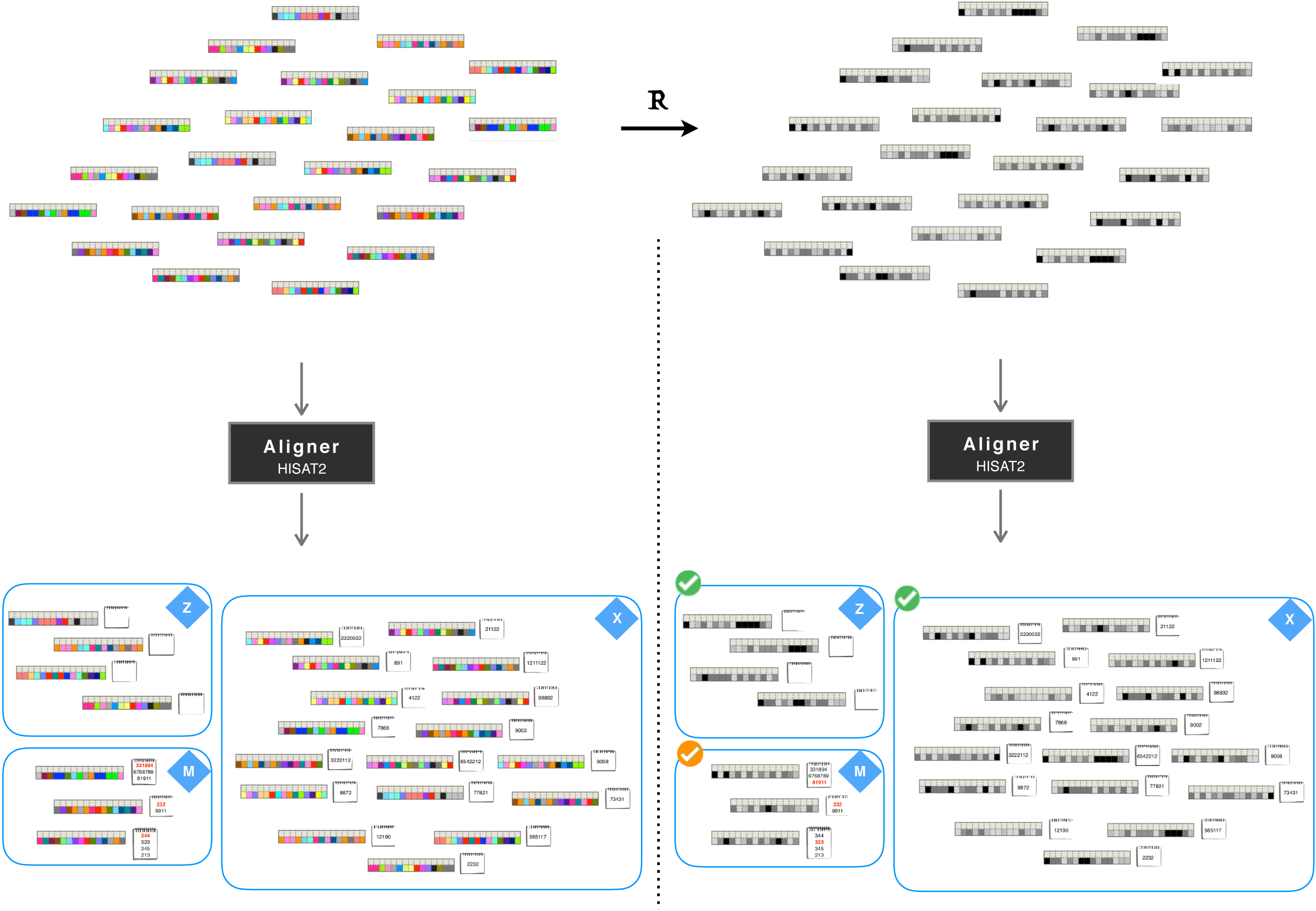
Schematic comparing the effect of rebinned reads on alignment.

## Discussion

We investigated the penalty functions that drive the alignment score system for read sequence alignment in a well-known quality-aware aligner. We then derived a simplification in the assignment of penalty values that reduces quality score scale granularity while keeping alignment scores unaffected. Consequently, this coarser quality score scale reduces storage footprint of sequence files with the advantage of entirely preserving read mapping percentages. In other words, we distorted quality scores without collateral impact on alignment.

The aligner in question was HISAT2, the modern version of the popular aligner Bowtie2, suitable for mapping genome and exome sequence data. Compared to other quality-aware aligners like Novoalign, HISAT2s approach to alignment score computation is straight-forward and deterministic, making it a good candidate to explore the relation and effect of quality scores and sequence alignment.

Simplifying the representation of quality scores is arguably a natural choice in the face of the sequence data explosion, and computational methods that approach the problem introduce collateral errors that are difficult to quantify. The assessment of quality score distortion has been attempted in some application domains (*17, 19, 37*) without clear consensus on the limits of safe lossy distortion levels. Meanwhile the increasing complexity of genomic assays, datasets and computational methods only adds to the difficulty of its potential quantification.

Nevertheless, even uniform requantization of the quality scores is a suitable approximation for high accuracy applications (*38*), and we have shown that this approach can be extended further to rebin coarsely quality scores without impact in sequence alignment.

In the light of the fast-paced sequencing technology progress, the utility of quality scores is at stake, as they are arguably unnecessary for many omics applications. We must therefore advocate for a feasible and pertinent granularity that suits each host application.

## Acknowledgments

This work was generously supported by the Swiss National Science Foundation.

## Footnotes

a https://www.ncbi.nlm.nih.gov/genbank/statistics/

b https://ccb.jhu.edu/software/hisat2/index.shtml

## References

1. T. S. Hey T, T. K, Microsoft Research (2001).

2. R. Kitchin, Big Data & Society (2014).

3. K. Wetterstrand, DNA Sequencing Costs: Data from the NHGRI Genome Sequencing Program (GSP), [ONLINE] Available at:https://www.genome.gov/about-genomics/fact-sheets/DNA-Sequencing-Costs-Data. [Accessed July 2, 2019].

4. Z. D. Stephens, et al., PLOS Biology (2015).

5. Genomics England. the 100,000 Genomes Project, [ONLINE] Available at: https://www.genomicsengland.co.uk/about-genomics-england/the-100000-genomes-project/ (2014). [Accessed June 22, 2019].

6. E. Birney, J. Vamathevan, P. Goodhand, bioRxiv (2017).

7. Next in the Genomics Revolution: The Era of the Social Genome | Veritas Genetics, [ONLINE] Available at: https://www.veritasgenetics.com/next-genomics-revolution-era-social-genome. [Accessed June 22, 2019].

8. J. Davis-Turak, et al., Expert Review of Molecular Diagnostics (2017).

9. M. Hosseini, D. Pratas, A. Pinho, Information (2016).

10. I. Numanagić, et al., Nature Methods (2016).

11. D. C. Jones, W. L. Ruzzo, X. Peng, M. G. Katze, Nucleic Acids Research (2012).

12. J. K. Bonfield, M. V. Mahoney, PLoS ONE (2013).

13. Å. Roguski, I. Ochoa, M. Hernaez, S. Deorowicz, Bioinformatics (2018).

14. A. El Allali, Source Code for Biology and Medicine (2019).

15. M. Hsi-Yang Fritz, R. Leinonen, G. Cochrane, E. Birney, Genome Research (2011).

16. F. Hach, I. Numanagić, C. Alkan, S. C. Sahinalp, Bioinformatics (2012).

17. I. Ochoa, M. Hernaez, R. Goldfeder, T. Weissman, E. Ashley, Briefings in Bioinformatics (2016).

18. D. L. Greenfield, O. Stegle, A. Rrustemi, Bioinformatics (2016).

19. A. A. Hernandez-Lopez, J. Voges, C. Alberti, M. Mattavelli, J. Ostermann, 2018 Data Compression Conference (IEEE, Snowbird, UT, 2018).

20. Illumina. NovaSeq 6000 System Quality Scores and RTA3 Software, Tech. rep. (2017).

21. H. Li, N. Homer, Briefings in Bioinformatics (2010).

22. N. A. Fonseca, J. Rung, A. Brazma, J. C. Marioni, Bioinformatics (2012).

23. G. Baruzzo, et al., Nature Methods (2017).

24. A. Hatem, D. Bozdaǧ, A. E. Toland, Ü. V.alyürek, BMC Bioinformatics (2013).

25. S. Thankaswamy-Kosalai, P. Sen, I. Nookaew, Genomics (2017).

26. A. D. Smith, Z. Xuan, M. Q. Zhang, BMC Bioinformatics (2008).

27. H. Li, J. Ruan, R. Durbin, Genome Research (2008).

28. H. Li, R. Durbin, Bioinformatics (2009).

29. Novocraft, Novoalign & NovoalignCS Reference Manual, Tech. rep. (2017).

30. B. Langmead, S. L. Salzberg, Nature Methods (2012).

31. H. Li, arXiv:1303.3997 [q-bio] (2013). 1303.3997.

32. R. Cánovas, A. Moffat, A. Turpin, Bioinformatics (2014).

33. G. Malysa, et al., Bioinformatics (2015).

34. Y. W. Yu, D. Yorukoglu, J. Peng, B. Berger, Nature Biotechnology (2015).

35. European Nucleotide Archive. T16M Metastatic liver tumor; File: SRR089705.fastq.gz, [ONLINE] Available at: https://www.ebi.ac.uk/ena/data/view/SRR089705.

36. European Nucleotide Archive. Gene expression data in skin fibroblast cells; File: SRR7093809.fastq.gz, [ONLINE] Available at: https://www.ebi.ac.uk/ena/data/view/PRJNA454681.

37. C. Alberti, et al., 2016 Data Compression Conference (DCC) (IEEE, Snowbird, UT, USA, 2016).

38. Illumina white paper. Reducing Whole-Genome Data Storage Footprint., Tech. rep. (2012).

